# ssHMM: Extracting intuitive sequence-structure motifs from high-throughput RNA-binding protein data

**DOI:** 10.1101/076034

**Authors:** David Heller, Martin Vingron, Ralf Krestel, Uwe Ohler, Annalisa Marsico

## Abstract

RNA-binding proteins (RBPs) play important roles in RNA post-transcriptional regulation and recognize target RNAs via sequence-structure motifs. To which extent RNA structure influences protein binding in the presence or absence of a sequence motif is still poorly understood. Existing RNA motif finders which produce informative motifs and simultaneously capture the relationship between primary sequence and different RNA secondary structures are missing. We developed ssHMM, an RNA motif finder that combines a hidden Markov model (HMM) with Gibbs sampling to learn the joint sequence and structure binding preferences of RBPs from high-throughput data, such as CLIP-Seq sequences, and visualizes them as a graph. Evaluations on synthetic data showed that ssHMM reliably recovers fuzzy sequence motifs in 80 to 100% of the cases. It produces motifs with higher information content than existing tools and is faster than other methods on large datasets. Examples of new sequence-structure motifs identified by ssHMM for uncharacterized RBPs are also discussed. ssHMM is freely available on Github at https://github.molgen.mpg.de/heller/ssHMM.

## 1 Background

RNA-binding proteins (RBPs), a class of proteins capable to bind RNA molecules, play a vital role in post-transcriptional control through processes such as RNA localization, RNA editing, RNA stability and splicing [2]. In human cells, hundreds of RBPs have been discovered but the detailed functional mechanisms of only a few have been explored so far [3, 2]. RBPs are known to recognize RNA molecules by their nucleotide sequence as well as their three-dimensional structure. Moreover, it has been found that many RBPs prefer binding to RNAs in specific structural contexts (Fig. 1) [1, 4]. To characterize the function of an RBP, it is crucial to first identify its interaction partners, i.e. the regulated gene transcripts. In most cases, the RNA targets of an RBP share at least one common local sequence or structure preference — a so-called motif, which fits into the binding pocket of the protein and thus facilitates the recognition of the RNA by the protein.

**Figure 1:**
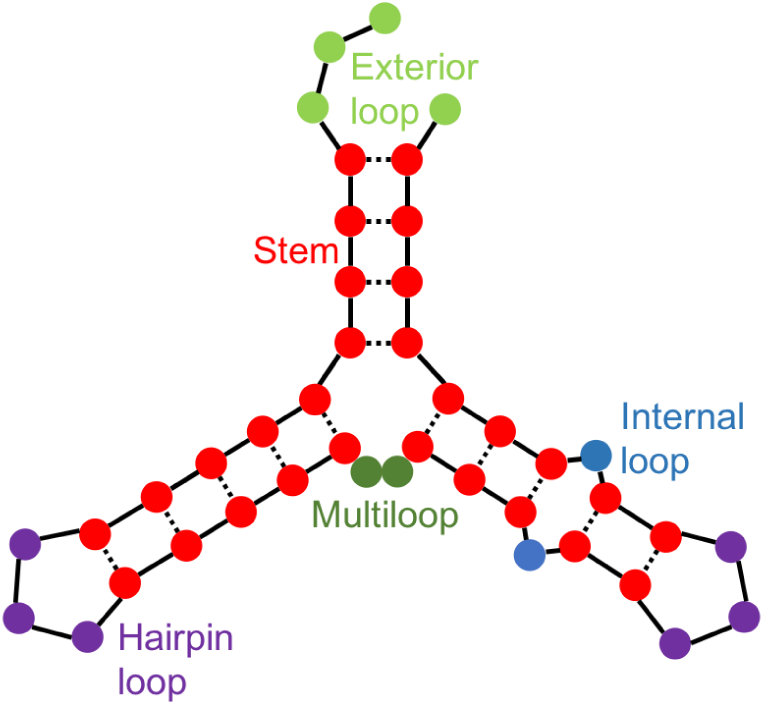
*Visual representation of five structural contexts of RNA - Colored nucleotides form the RNA strand (solid line) and some of them are bound together by base-pairing (dashed lines). The five structural contexts are represented by five colors: stems (red), exterior loops (light green), hairpin loops (purple), internal loops (blue), and multiloops (dark green). Figure adapted from [1]*.

Several approaches for motif finding, i.e. computationally extracting an unknown motif from a set of target sequences, have been developed for transcription factor (TF) binding sites in DNA sequences.

They can be categorized into four major classes [5]: (1) Enumerative algorithms, which count the occurrences of exact *k*-mers in the sequence set to find over-represented words [6, 7, 8]. (2) Optimization algorithms based on expectation maximization (EM) to simultaneously optimize a position weight matrix (PWM) description of a motif [9] and probabilities of motif starts in the associated sequences. A popular implementation of the EM algorithm is the *MEME* software [10]. (3) Algorithms based on probabilistic optimization, such as Gibbs sampling, which iteratively sample from the conditional distribution of one motif start at a time [11].(4) Affinity-based (motif) models which parametrize and fit a function representing the binding affinity of a protein (e.g. a TF) for a set of words [12, 13, 14].

Modern high-throughput methods for the analysis of protein binding interactions yield thousands of RNA or DNA binding sites. Compared to experimental methods for detecting TF binding sites on DNA, high-throughput protocols for protein-RNA interactions are relatively new. Among them, in-vitro evolutionary methods, such as SELEX [15] and RNA-compete [16], which identify high-affinity RNA ligands within pools of randomly or specifically selected sequences. Alternatively, various crosslinking and immunoprecipitation (CLIP) methods have been introduced [17, 18, 19], which rely on covalent crosslinking of an RBP to its RNA target in living cells, followed by isolation of RBP-RNA fragments and deep sequencing. The RNA sequences (reads) produced by CLIP-Seq protocols can be mapped back to the genome, and peak calling tools, such as Piranha [20] or PARalyzer [21], can be used to identify high-fidelity RBP binding sites from the read levels [21, 20, 22].

Although much work has been done in the area of DNA motif finding, few approaches have been developed for RNA motifs (Table 1). One of the reasons for this is that high-throughput methods for RBP binding site profiling, such as CLIP-Seq, are relatively new. Additionally, the binding of RNA depends not only on the RNA’s nucleotide sequence but also on its three-dimensional structure. Consequently, motif finders that work well for finding motifs in linear DNA cannot easily be applied to RNA, but have to be extended to take the RNA secondary structure into account. This is a difficult task due to the noisy nature of computational RNA secondary structure prediction, the fuzzy patterns of sequence-structure motifs, the large set of input sequences where the motif could possibly be contained, and the potentially large number of false positives among the RBP binding sites called from CLIP-Seq experiments [23].

**Table 1:**
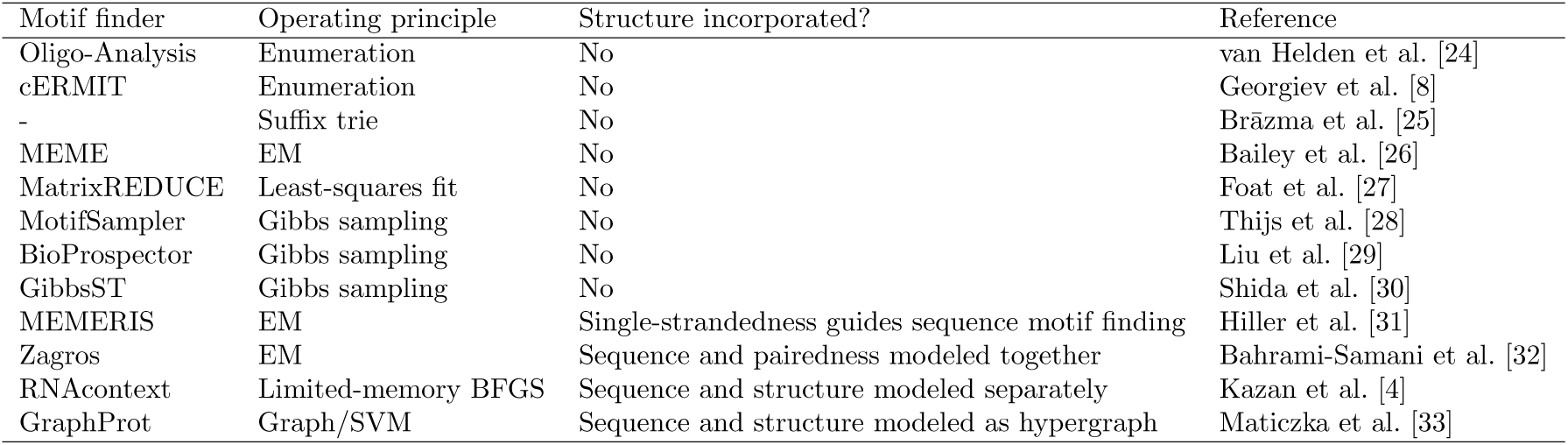
*A selection of RNA motif finding algorithms. The third column indicates whether the motif finder incorporates RNA secondary structure in any way.*

Existing RNA motif finders either address only part of the problem or employ machine learning models which are harder to interpret in terms of sequence-structure preferences (Table 1). The first tool to incorporate RNA secondary structure into motif prediction was *MEMERIS* [31], an extension of the *MEME* EM algorithm. It uses single-strandedness information as a prior to guide motif finding to single-stranded regions based on the assumption that most RBPs prefer to bind in single-stranded regions. However, recent studies have shown that several RBPs bind to stem-like regions, rather than to single-stranded loops [1]. Therefore, the main limitation of *MEMERIS* is that it misses binding motifs for proteins with stem-loop preference. Moreover, it does not take into account the full spectrum of RNA structures and does not distinguish between different single-stranded contexts, such as hairpin loop and internal loop. Another extension to the *MEME* software is *Zagros* introduced by Bahrami-Samani et al. [32], where the authors account for secondary structure (paired or unpaired only) and crosslinking modifications in the EM framework.

Several methods exist to incorporate the full secondary structure into RNA motif finding. In 2010, Kazan et al. introduced *RNAcontext*, an affinity-based model which learns both sequence and structure preferences of an RBP, considering several structural contexts [4]. In *RNAcontext*, each nucleotide in the input sequence is annotated with a distribution over four structural contexts. *RNAcontext* learns the RBP binding affinity and optimizes the model’s parameters from both sequence k-*mers* and structural profiles. Affinity values are obtained by experimental assays such as *RNA-compete*, or can be set to discrete classes, e.g. *bound* and *unbound* from CLIP-Seq experiments. *RNAcontext* outperformed *MEMERIS* in classifying bound versus non-bound RNA sequences on a selection of nine proteins, but its performance on data other than *RNAcompete* assays is disputable.

*GraphProt* by Maticzka et al. uses graph kernel-based support vector machines (SVM) trained on a large number of features from a hypergraph to learn sequence and structure preferences of RBPs [33] from high-throughput data. *GraphProt* produces a motif visualization in the form of separate sequence and structure logos. Feature interpretation from the hypergraph SVM model is not straightforward and the two logos are indirectly generated from the top-scoring k-*mer* nucleotide sequences and structure profiles. Alternatively, the trained model can be used to predict novel binding sites in the same organism. Like *RNAcontext*, *GraphProt* requires at least a positive and a negative input dataset for training. In a binary classification setting it was shown to be superior to *RNAcontext* for a large set of RBPs.

While *RNAcontext* and *GraphProt* are designed to accurately distinguish bound from unbound sites, a tool for *de novo* identification of sequence-structure motifs from RBP-bound sequences is missing.

In this paper, we propose ssHMM, a novel tool to identify *de novo* sequence-structure motifs in a set of RNA sequences bound by a certain RBP. Our method, trained on CLIP-Seq experimental data, and based on hidden Markov models to represent both sequence and structure preferences has several advantages compared to previous approaches: (1) It identifies a combined sequence-structure motif which characterizes the unique features of the binding site rather than outputting two separate logos for sequence and structure. (2) It models a spectrum of five different structural contexts (stem and four different single-stranded loop contexts). (3) It makes use of Gibbs sampling and sub-optimal RNA shape predictions, which might be closer to the native RNA conformation, to optimize the sequence-structure pattern. (4) It is designed with the purpose of producing an interpretable motif model which can be intuitively visualized and easily understood. ssHMM outperformed both *RNAcontext* and *GraphProt* in retrieving the correct sequence motif from synthetic data while being comparable to *MEMERIS*. On the other hand, when applied to real biological data from CLIP-Seq experiments, our method produced more informative motifs than *RNAcontext* and *GraphProt* for most of the analyzed RBPs and proved to be considerably faster than *MEMERIS* on large datasets. A global analysis of our results reveals that the structure preference of an RBP decreases with increasing strength of the sequence motif. We also demonstrate the ability of our tool to reveal new insights into previously uncharacterized RBP motifs and their biological implications for RBP function. For instance, we describe the sequence-structure preferences of RBPs involved in small RNA processing, such as DGCR8 and DICER, as well as YY1, a transcription factor which has recently been shown to bind RNA as well as DNA [34]. ssHMM is freely available for download on Github (https://github.molgen.mpg.de/heller/ssHMM), is easy to use, and can be applied to characterize novel motifs in any set of input RNA sequences.

## 2 Results

Here, we present ssHMM, a *de novo* motif discovery tool which combines hidden Markov models (HMMs) with Gibbs sampling to learn the joint sequence and structure binding preferences of an RBP. ssHMM is trained on high-throughput RNA-binding protein data from CLIP-Seq experiments or any other experimental protocol yielding large numbers of RNA sequences. After training, the resulting model can be visualized as an intuitive graph logo.

### 2.1 The ssHMM model

The sequence-structure hidden Markov model (ssHMM) constitutes the core of the motif finder presented here. It is trained on two types of “sequences” — the actual RNA nucleotide sequences corresponding to the RBP binding sites as detected by means of high-throughput experiments and their corresponding RNA structures (Fig. 2). The latter are encoded as sequences of symbols representing different structural contexts. RNA structures are predicted from the primary sequences using either *RNAstructure* or *RNAshapes* [35, 36]. The user can choose which of these two structure prediction tools to use. Each letter in an RNA structure sequence has to denote the kind of secondary structure that the corresponding nucleotide in the primary sequences assumes albeit abstracting from the predicted base pairing. More details about the encoding of predicted structures into symbols of structural contexts are given in the Methods Section. All results presented here were generated using structure predictions from *RNAshapes*.

**Figure 2:**
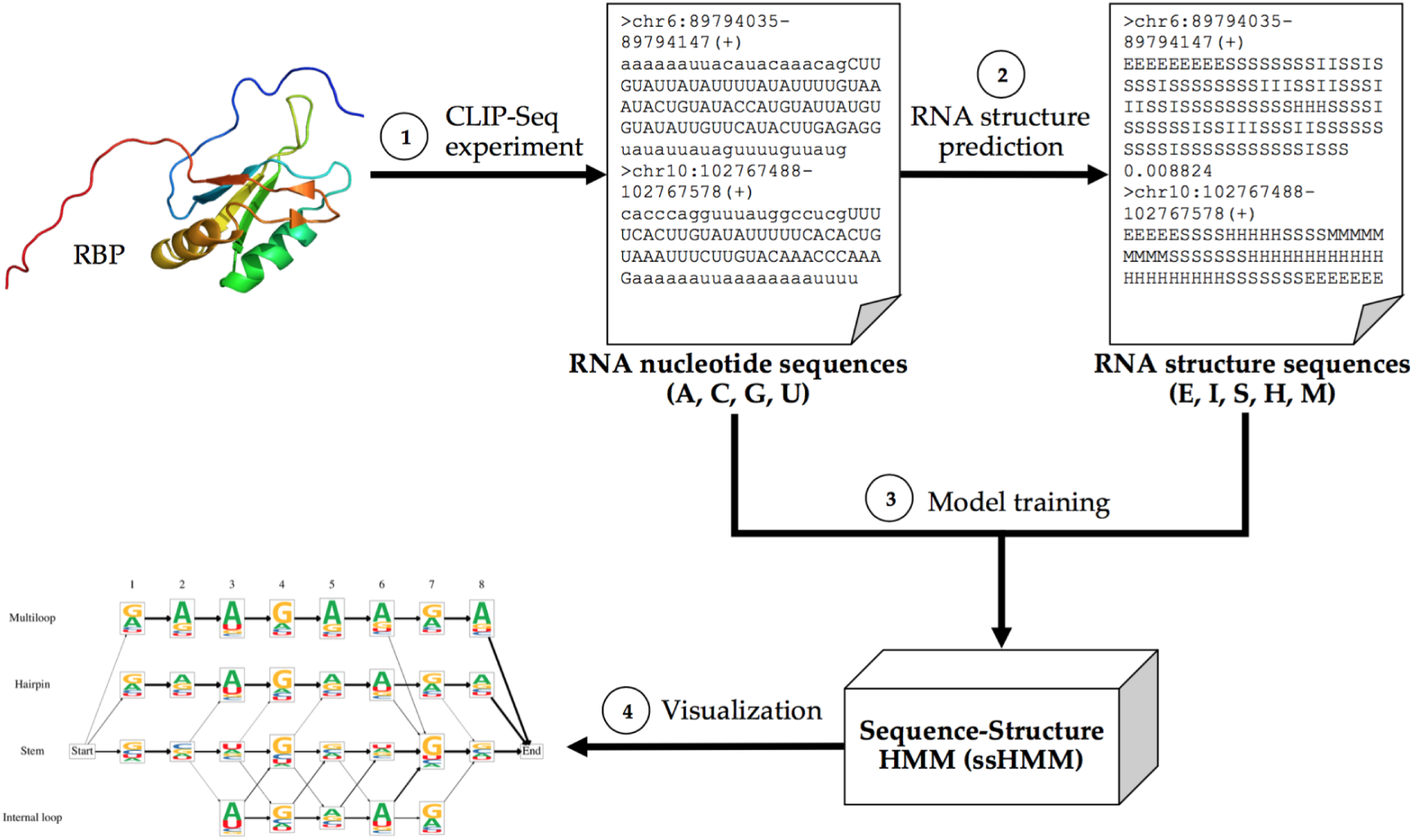
*Overview of the motif finder workflow - (1) A CLIP-Seq experiment yields nucleotide sequences of RNAs (in uppercase) that were putatively bound by a specific RBP. The genomic region surrounding the binding site was added in lowercase to benefit the next step. (2) The most likely structural conformations of each nucleotide sequence are computed by a structure prediction tool* (RNAshapes *and* RNAstructure). *(3) The ssHMM is trained with the nucleotide and structure sequences. (4) The final ssHMM can be visualized in an intuitive way similar to a sequence logo*.

In contrast to previous approaches, ssHMM simultaneously captures the sequence and structure binding preferences of an RBP. This is reflected in the HMM topology: The states of the model represent five different structural contexts: stem, hairpin loop, multiloop, internal loop, and exterior loop (Fig. 1), while its emissions represent the four RNA nucleotides. The rationale behind this topology is that RBPs might recognize their RNA targets by both their nucleotide sequence and their structure. ssHMM not only captures sequence and structure preferences of an RBP, but also models the relationship between RNA sequence and structure at RBP binding sites, which helps to elucidate whether a specific structural context is required or not in order to observe a certain sequence motif.

### 2.2 Training the ssHMM with Gibbs sampling

The sequence-structure HMM is trained with a Gibbs sampling approach to estimate two sets of required variables: Firstly, the position of the motif in each long RNA input sequence has to be determined. Secondly, the optimal RNA structure among several possible conformations produced by the structure prediction tool needs to be estimated. Our Gibbs sampling method enables the simultaneous estimation of the HMM parameters and both variable sets.

At the outset, initial values for the two sets of variables are set. Then, an iterative optimization process begins that alternates between reestimating the ssHMM and the unknown variables. Every iteration consists of three steps: In the first step, one of the input sequences is randomly chosen and the ssHMM is re-estimated using all but the chosen sequence. In the second step, the motif position and the optimal RNA structure of the chosen sequence are reestimated. In the third step, a new motif position and optimal RNA structure are estimated for the chosen sequence. The algorithm terminates when the improvement per iteration in the likelihood of the data drops below a user-defined threshold. At this point the algorithm should have found parameter values such that the likelihood of the ssHMM given the input data is close to optimal. To accelerate the training and obtain better results, we pick initial values for the Gibbs sampler by training a separate sequence-only HMM using the Baum-Welch algorithm. Alternatively, the initial values can be chosen randomly. For a detailed description of the model training and the model parameters see the Methods Section and Additional File 1.

### 2.3 ssHMM visualization

The trained HMM can be visualized as a graph in which each state is represented by one node. Similar to a sequence logo, the nodes of the graph visualize the emission preferences of the corresponding HMM state with stacks of colored nucleotide letters. These stacks indicate which bases are prevalent at each binding site position in each structural context. The transition probabilities between the HMM states are visualized as arrows. The thicker an arrow between two states, the more likely is a transition between the two. Arrows corresponding to transition probabilities lower than 5% are not displayed to increase clarity.

### 2.4 Evaluation of ssHMM on both synthetic and biological data

We evaluated the performance of ssHMM on synthetic and biological datasets and compared it with three state-of-the-art approaches for RBP sequence-structure motif finding: *MEMERIS*, *RNAcontext*, and *GraphProt* [31, 4, 33]. The synthetic datasets were randomly generated with specific characteristics, while the biological datasets comprise several CLIP-seq experiments publicly available from various biological databases and repositories. More details about the used datasets and their sources can be found in the Methods Section. In the following, we describe the results of our analyses.

### 2.5 ssHMM reliably recovers sequence motifs from synthetic sequences

The evaluation of a motif finder with synthetic datasets yields several advantages. In contrast to biological sequences, synthetic sequences are generated specifically to contain a certain motif and a certain amount of noise. As the correct motif, that is supposed to be recovered by the motif finder, is known beforehand, the motif finding performance can be objectively evaluated. It can be explored how well the motif finder extracts the hidden motif and which sequence properties influence motif finding performance.

All three state-of-the-art tools use different structure representations so that the produced structure motifs cannot be compared directly. *RNAcontext*, for instance, produces only a single set of relative structural context affinities over the entire motif. *MEMERIS* does not output a structure motif at all and regards only the propensity of a nucleotide to be single-stranded. Therefore, we confined our analysis to an evaluation of how well the tools recover implanted sequence motifs in hairpin loop regions from synthetic datasets.

To produce the datasets, we implanted fuzzy sequence motifs into background sequences at random locations. 12 datasets were generated which varied in motif information content (1.0 and 0.5), background sequence type (random and 3’UTR), and fraction of motifs in a hairpin loop. After running the tools, Kullback-Leibler (KL) divergence between the recovered sequence motif and the originally implanted motif was calculated. Thus, the KL divergence can serve as a measure of how well the tools recover a motif hidden in sequences. For each dataset and tool, three repetitions were performed and the mean recovery rate was reported. In the Methods Section we explain the evaluation techniques in more detail.

Our analysis confirms that ssHMM, together with *MEMERIS*, is able to perfectly recover 100% of the motifs with information content 1.0 (Fig. 3). For these rather clearly defined motifs *RNAcontext* reached results considerably below 100%.

**Figure 3:**
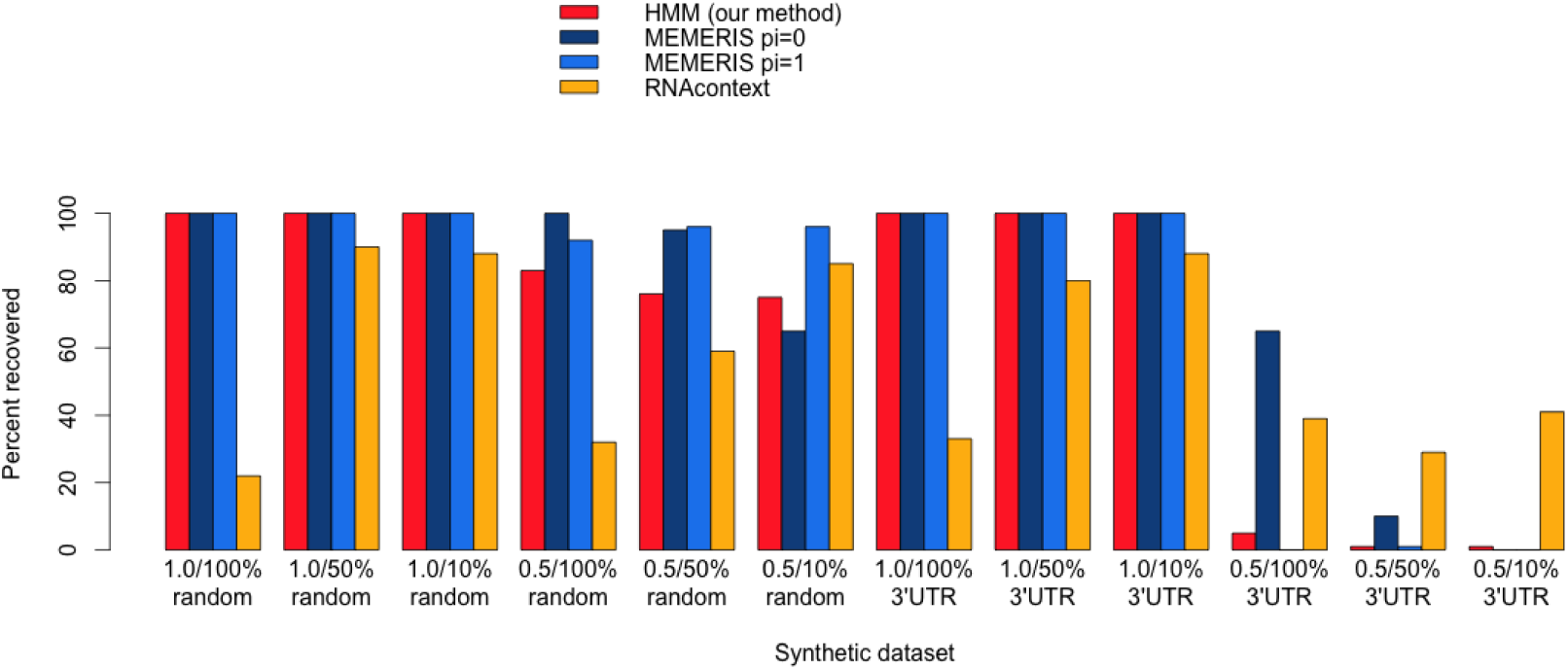
*Comparison of recovery rates of ssHMM*, MEMERIS, *and* RNAcontext *on synthetic data - Shown is the percentage of successfully recovered motifs from 12 synthetic datasets. The colors represent the different tools. For* MEMERIS, *two different values for the pi parameter are plotted. The pi parameter determines the importance of the secondary structure relative to the sequence. The labels on the x axis show the motif information content, hairpin loop fraction, and background sequence type of each dataset*.

*GraphProt* displayed a very weak performance over all synthetic datasets (data not shown). Due to the synthetic nature of the sequence-structure motifs considered here, sequences might not form proper and biologically valid secondary structures and cannot be represented in a valid graph structure. *GraphProt* heavily depends on a graph to extract its model features, rather than on short sequence motifs. Therefore, a comparison of our tool with *GraphProt* in this test setting would not be completely fair, which is why *GraphProt* is not included here.

The recovery of fuzzy motifs with an information content of 0.5 depends highly on the type of background sequences they were implanted in. For random background sequences, ssHMM reached slightly lower recovery rates than *MEMERIS* but higher rates than *RNA-context*. For 3’UTR background sequences however, all tested motif finders had great difficulties and reached recovery rates well below those for random background sequences. 3’UTRs contain strong confounding regulatory sequence signals important for RNA post-transcriptional regulation other than the implanted motif. That makes the task of retrieving such a weak synthetic motif much harder.

Overall, ssHMM achieved results that were superior to *RNAcontext* and comparable to *MEMERIS*. The mean sequence recovery rates for ssHMM, *RNAcontext*, *MEMERIS* (pi=0), and *MEMERIS* (pi=1) were 70.08%, 57.17%, 77.92%, and 73.75%, respectively.

### 2.6 ssHMM can distinguish between real binding sites and background

With our motif finder we analyzed 21 different PAR-CLIP, HITS-CLIP, and iCLIP datasets for 18 different RBPs from various sources. With the exception of two mouse datasets, all datasets stemmed from human HEK293 and HeLa cells.

To confirm that our motif finder is able to distinguish between real binding sites and background sites, we analyzed the difference in log-likelihood between a positive and a negative test set given a trained motif model. In this analysis the Wilcoxon rank sum test yielded p-values smaller than 0.01 for all CLIP-Seq datasets (Additional File 1). For the majority of proteins, the p-value was smaller than 2.2e-16. The significance in log-likelihood differences demonstrates that our trained motif model can distinguish between real binding sites and background sites.

### 2.7 Motifs identified by ssHMM possess high information content (IC)

Comparing the performance of different motif finders on CLIP-Seq data proves difficult. The reason is that some approaches, including our motif finder, use a generative model which is trained solely on positive data (i.e. binding sites from CLIP-Seq experiments), unlike *RNA-context* and *GraphProt* which are trained also on negative data. While the parameters of a generative model are optimized to describe the positive data best, discriminative models are trained to distinguish positive from negative data. Due to these differences a fair comparison of our tool with other motif finders requires special care.

Moreover, the primary goal of motif finding is not the prediction or classification of binding sites but the retrieval of an informative binding site motif. Therefore, we are convinced that the recovered motif’s information content is an appropriate evaluation measure.

We computed three variants of the motif information content on three different alphabets *A*:

- Information content of the sequence motif (*A* = {*A, C, G, U*})
- Information content of the structural motif (if applicable, *A* = {*E, I, S, H, M*})
- Information content of sequence and structure combined (if applicable, *A* = {*A, C, G, U*} × {*E, I, S, H, M*})

We compared the information content of ssHMM with that of the two other sequence-structure motif finders *RNAcontext* and *GraphProt*. *MEMERIS*, which yields only a sequence motif, was not considered for this comparison. To ensure a fair comparison of the information content, we distinguish two ways of computing it: For comparison with *GraphProt*, we obtained the information content from the top-scoring 1000 sequences, as this is the methodology that GraphProt uses to produce its motif logos. For comparison with *RNAcontext*, we computed the information content directly from the trained model. The calculation of information content is explained in greater detail in Additional File 1.

Our motif finder produced more expressive motifs than *GraphProt* and *RNAcontext* for almost all proteins (Fig. 4 (a–c)). The mean information content of all 21 sequence motifs (from the top 1000 sequences) was 1.23 and 0.98 for ssHMM and *GraphProt*, respectively. For all 21 structure motifs, the mean information content was 1.2 and 0.8, respectively. Only for five of the proteins did *GraphProt* produce a more expressive structure motif than ssHMM. One explanation for these cases could be that some proteins need a larger structural context to define their preferences, in which case *GraphProt* benefits from the graph representation. By combining the sequence and structure alphabets one can also obtain sequence-structure motifs. For all proteins, the combined motifs by our motif finder had a higher information content than those by *GraphProt*. Their mean information content was 2.57 and 1.73 for ssHMM and *GraphProt*, respectively.

**Figure 4:**
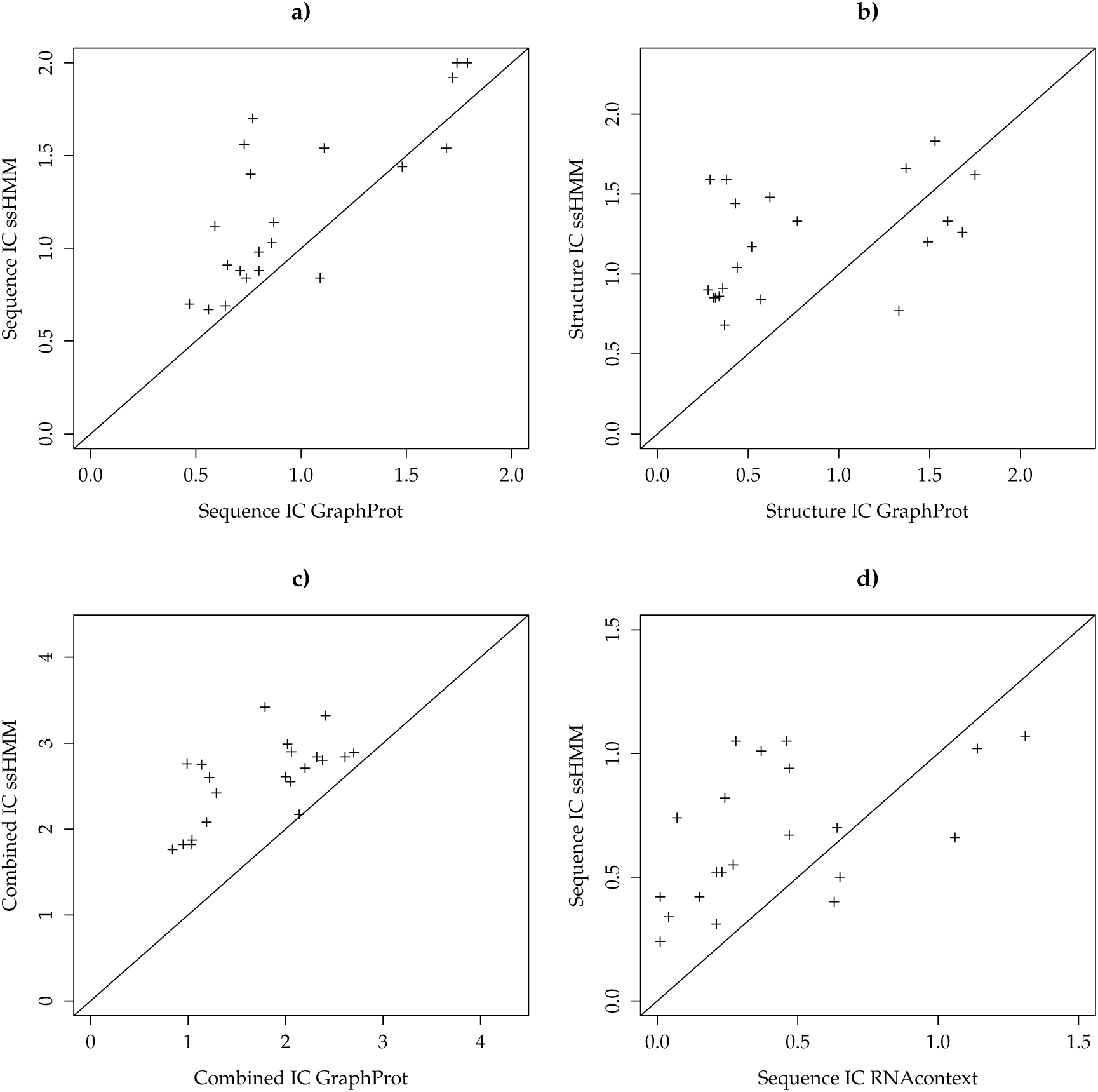
*ssHMM produces motifs with a higher information content than* GraphProt *and* RNAcontext - ***(a-c)** The information content of a) sequence motifs, b) structure motifs, and c) sequence-structure motifs was compared for* GraphProt *(x-axis) and ssHMM (y-axis). Points above the solid line represent protein datasets for which ssHMM yielded a larger information content than* GraphProt. *For points below the line,* GraphProt *produced the motif with higher information content. **(d)** The information content of sequence motifs produced by* RNAcontext *(on the x-axis) was compared to that of motifs by ssHMM (y-axis). Points above the solid line represent a larger information content for ssHMM whereas points below the line denote larger information content for* RNAcontext.

The motifs obtained directly from ssHMM were generally more expressive than those by *RNAcontext* (Fig. 4d,). On average, ssHMM and *RNAcontext* sequence motifs (directly from the model) had an information content of 0.66 and 0.42, respectively. Only for five of the proteins did *RNAcontext* produce a more expressive motif than ssHMM. For three of those (DGCR8, FXR2, and PUM2), the difference was larger than 0.2 due to the fact that the motif recovered by *RNAcontext* was different from the one retrieved by both *MEMERIS* and ssHMM.

### 2.8 Negative correlation between sequence and structure specificity in RBP binding sites

We investigated more in detail the relationship between sequence and structure in RBP binding sites. Over all CLIP-Seq datasets, a significant negative correlation of −0.536 (Spearman’s rank correlation, p-value 0.006) can be observed between sequence and structure information content (Fig. 5).This observation is concordant to what Schneider et al. described in 1986: Binding sites tend to contain approximately as much information as is necessary for them to be recognized [37]. In RNA-RBP binding, sequence and structure specificity seem to complement each other so that RBPs with a strong sequence preference tend to exhibit only a weak or no structure preference and vice versa.

**Figure 5:**
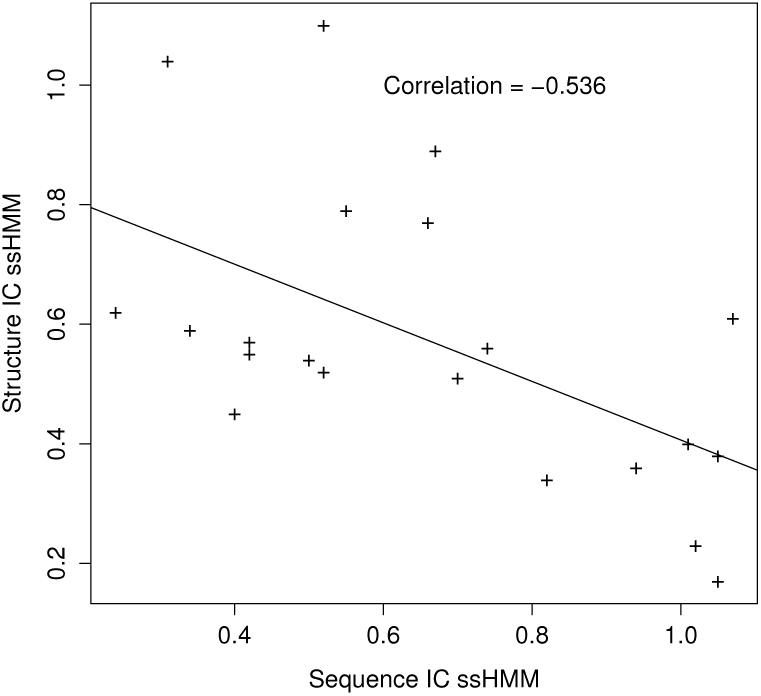
*Sequence and structure information content of motif models trained on CLIP-Seq datasets are negatively correlated. - Shown are the sequence information content (x-axis) and structure information content (y-axis) of ssHMM motifs for all 21 CLIP-Seq datasets. Both are negatively correlated with a significant Spearman’s rank correlation coefficient of −0.536 (p-value 0.006). Plotted is also the linear regression line.*

### 2.9 ssHMM recovers previously validated motifs from CLIPSeq data

Table 2 shows the output of our motif finder for five selected CLIP-Seq datasets. For the full list of results from all datasets, including the literature motifs, motifs recovered by *MEMERIS*, *RNAcontext*, and *GraphProt*, and their corresponding information contents, see Additional File 2. For three of the proteins, Nova, QKI, and DICER, motifs have been characterized before which confirms that ssHMM recovers correct motifs from biological data.

**Table 2:**
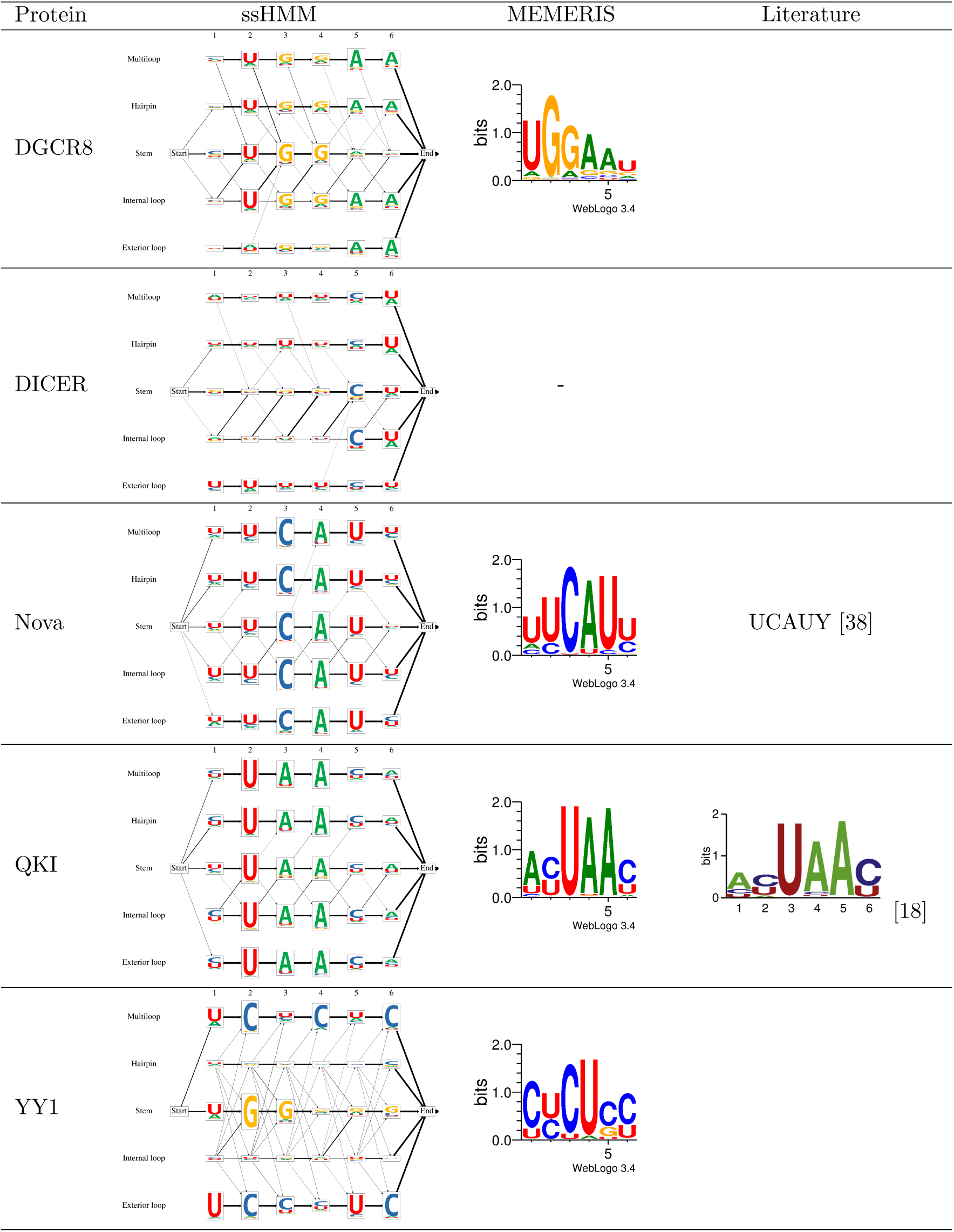
*Visualization of trained ssHMMs for five selected CLIP-Seq datasets. In the third column, the sequence logo recovered by MEMERIS is shown for comparison. On the DICER dataset,* MEMERIS’ *runtime exceeded 24 hours and was terminated. For some proteins, a literature motif from other studies is shown in the fourth column.*

| Protein | ssHMM | MEMERIS | Literature |
| --- | --- | --- | --- |
| DGCR8 |  |  |  |
| DICER |  | - |  |
| Nova |  |  | UCAUY [38] |
| QKI |  |  | [18] |
| YY1 |  |  |  |

#### 2.9.1 Nova

Nova is an RBP exclusively expressed in neurons within the central nervous system that is putatively involved in RNA alternative splicing and RNA metabolism. The CLIP-Seq dataset we analyzed was therefore obtained from mouse brain cells. From it, our motif finder recovered a clear UCAU sequence motif and U-rich flanking positions. This agrees with the motifs from *MEMERIS* and findings from other studies [38, 39, 1] which identified unstructured and U-rich regions as preferred binding environment for NOVA. The visualized ssHMM displays a considerable preference for the single-stranded multiloop and hairpin loop contexts.

#### 2.9.2 QKI

The Quaking homolog (QKI) is an RBP regulating pre-mRNA splicing, mRNA export, mRNA stability, and protein translation [40]. It is suspected to play a role in schizophrenia [41]. The ssHMM visualization for this protein shows a strong UAA sequence motif across all structural contexts. This is in accordance with the motifs retrieved by *MEMERIS*, *RNAcontext*, *GraphProt*, and other previous studies [18]. Our motif finder also found that QKI prefers to bind hairpin loops and, to a lesser degree, multiloops, exterior loops, and stem structures.

#### 2.9.3 DICER

The protein DICER was investigated in detail here because it is a key player in the microRNA biogenesis pathway. DICER is specifically involved in the processing of microRNA precursors (pre-miRNAs) into double-stranded RNA fragments which then give rise to the ~21-nt-long mature microRNAs. It is known that DICER binds double-stranded RNA structures and that this structural context determines its specificity [42, 43], while nucleotide sequence does not play a role in determining DICER efficiency [42]. ssHMM confirms previous observations about DICER’s binding preference for the stem context, the lack of a strong sequence binding motif and the preference for C nucleotides in the penultimate terminal stem position [42].

### 2.10 ssHMM recovers new motifs from CLIP-Seq data

With our motif finder, we were able to recover novel motifs from proteins with, until now, undescribed sequence-structure binding preference.

#### 2.10.1 DGCR8

Together with the Drosha protein, the RBP DGCR8 forms the so-called Microprocessor complex which is involved in one of the first steps of microRNA biogenesis and is responsible for recognizing and releasing pre-miRNA hairpins from large primary microRNA (pri-miRNA) transcripts. The two proteins in the complex have distinct and complementary tasks: DGCR8 recognizes and binds primary miRNAs while Drosha cuts them and thus converts them to pre-miRNAs [44]. The sequence-structure preference of DGCR8 is not very well known so far. Previous studies have explored the relationship between RNA-binding properties of DGCR8 and pri-miRNA processing efficiency and found that DGCR8 binds both double-stranded and single-stranded transcripts with similar affinity, indicating a poor specificity of DGCR8 for a specific structural context [45].

Our analysis on a DGCR8 PAR-CLIP dataset revealed a slightly different scenario: ssHMM recovered a UGGAA sequence motif. This motif was identically retrieved by *MEMERIS* and, in more fuzzy form, by *GraphProt*. A remarkable observation can be made when comparing the sequence motifs from the different structural contexts based on ssHMM: While the single-stranded hairpin and internal loop contexts display the full UGGAA sequence motif, the stem context seems to miss the last two bases. It only exhibits the shortened motif UGG. ssHMM also reflects a strong preference for the stem context which is in accordance with the structural motif found by *GraphProt* and the fact that DGCR8 contains two double-stranded RNA-binding domains [46, 44]. This is in line with the findings from two previous studies [47, 48] aimed at characterizing sequence-structure determinants of efficient pri-miRNA processing. These studies identified a basal, highly-conserved UG dinucleotide motif in a stem context at RNA positions −14 and −13 from the Drosha cleavage sites to be involved in enhanced pri-miRNA processing. Neither study provides, however, any reason for the molecular mechanisms behind the function of the UG motif. From our ssHMM analysis, we can suggest that the UG dinucleotide might be contributing to the specificity of DGCR8 binding in a double-stranded structural context.

#### 2.10.2 YY1

Recently, transcription factors (TFs) that bind both DNA and RNA have gained considerable attention thanks to their prominent role in RNA-mediated transcriptional regulation of gene expression. In particular, a recent study about the role of the TF YY1 in mouse embryonic stem cells showed that RNAs transcribed from regulatory elements such as promoters and enhancers contribute to stabilizing DNA occupancy of this transcription factor [34]. While the DNA binding motif of TF YY1 is known, its RNA specificity has not yet been investigated. We analyzed the YY1 CLIP-Seq dataset from [34] and we derived, for the first time, a potential sequence-structure RNA binding motif for YY1. ssHMM revealed two major preferred sequence-structure contexts for YY1: a strong CU-rich motif in a multiloop context and a G-rich stem motif (Table 2). Although MEMERIS was able to retrieve the CU-rich sequence motif, it was not able to reveal the stem motif, even when run with different sets of parameters and searching for multiple motifs simultaneously. In addition, *MEMERIS* was not able to determine the location of the CU-rich motif in the multiloop context, simply because it does not differentiate between different single-stranded structure types. Both *RNAcontext* and *GraphProt* recovered a motif which merges both sequence motifs recovered by ssHMM (Additional File 2). But while *GraphProt* located the motif in the stem, *RNAcontext* located it in a hairpin loop context. This difference in the structure preference might be due to the different structure prediction tools employed by *RNAcontext* (*SFOLD*) and *GraphProt*/*ssHMM* (*RNAshapes*). The identification of new RBP motifs like the one for YY1 can contribute to shedding light on RNA functional characterization. In addition, this example shows that our approach can retrieve more than one preferred sequence motif, can determine their respective structural contexts, and can help characterizing protein preferences where other approaches return ambiguous results.

### 2.11 Motifs recovered from CLIPSeq datasets are independent of chosen structure prediction tool

Although tools for RNA structure prediction often yield differing results, we could show that the results of our motif finder are independent of the chosen structure prediction tool. For all 21 protein datasets, we compared the motifs recovered by ssHMM using structures predicted either by *RNAshapes* or *RNAstructure*. We observed that the two motifs were highly similar for all proteins (Additional File 3). Consequently, the motif finding results are robust to the choice of a particular structure prediction tool (*RNAshapes* or *RNAstructure*).

### 2.12 ssHMM can efficiently handle large datasets

In order to be suitable for scientific use, modern motif finders are required to process large datasets in reasonable time. CLIP-Seq is a high-throughput protocol and produces datasets that commonly hold tens of thousands of RBP binding sites. It is therefore vital that motif finding algorithms scale well with input datasets of increasing size.

To assess the runtimes of *GraphProt*, *RNAcontext*, *MEMERIS* and ssHMM, we took runtime measurements on datasets of increasing size. The four different motif finders did not scale equally well on the datasets (Additional File 1). The runtimes of both *GraphProt* and *RNAcontext* increased linearly with the input size. But while *GraphProt* exhibited a runtime comparable to ssHMM, *RNAcontext* took more than 10 times as long as the others on all datasets and its runtime increased also much more steeply. This was mainly due to the secondary structure prediction algorithm *Sfold* that *RNAcontext* employs. Its share of *RNAcontext*’s runtime amounted to more than 90%. When regarding only the training of the model, *RNAcontext* displayed a runtime progression similar to *GraphProt* and ssHMM.

For *MEMERIS*, we observed a quadratic runtime progression. While it was the fastest of the four approaches on the smallest dataset tested, it was overtaken by ssHMM and *GraphProt* on the largest. Although the largest dataset only comprised five times as many sequences as the smallest, *MEMERIS*’ runtime increased by a factor of 34.9.

Our motif finder seems to scale gently quadratically with increasing input size although the measurements are hard to generalize. As the algorithm involves randomness, the runtimes exhibited a much higher variance than those of the other tools. Gibbs sampling performs a random walk in an unknown probability landscape so that a theoretical complexity analysis is hard. Generally, the runtime of ssHMM increased slower than the input size. When comparing 200 to 1000 sequences, the runtime increased only by a factor of 4.5. For the two largest datasets, our motif finder was the fastest of all approaches.

All in all, *GraphProt* and ssHMM seem to be the only methods able to process large datasets in reasonable time, while *MEMERIS*’ runtime increases quadratically and *RNAcontext* is inhibited by its slow structure prediction tool.

## 3 Discussion

Knowing the sequence-structure specificity of RNA-binding proteins is essential for understanding RNA post-transcriptional regulatory processes, in the same way as sequence specificity of DNA-binding factors is crucial to develop models of transcriptional gene regulation. While several tools have been developed to extract *de novo* sequence motifs from sets of DNA sequences, for RNA sequences the secondary structure around the binding site influences their function and activity [1, 49, 50]. We developed ssHMM, a *de novo* motif finder capable of extracting sequence-structure RNA binding motifs from large sets of RNA sequences generated from genome-wide experiments such as CLIP-Seq.

With this study, we address four main points. First, our approach shows the advantages of incorporating the full spectrum of RNA structures into the motif model. ssHMM is based on a hidden Markov model (HMM) which consists of states representing the binding context formed by the different RNA structures. It is therefore able to estimate the binding sequence and structure specificities simultaneously. In contrast to *MEMERIS* which distinguishes only double-stranded and unstructured regions, ssHMM takes five different structural contexts into account. This gives a detailed insight into the preferred structural contexts of an RBP. ssHMM can, for instance, uncover an RBP’s specific preference for multiloops or hairpin loops instead of single-stranded regions in general, as shown for YY1 in the Results Section. Additionally, ssHMM captures the structural preference for every individual position of the RBP binding site and even models transitions between the structural contexts within the binding site. *RNAcontext*, in contrast, outputs only the global structural preference of an RBP but not for individual binding site positions. *MEMERIS* not even produces a structure logo but a sequence motif only. The analysis of several CLIP-Seq datasets with such a combined sequence-structure model allowed us to reveal an interesting anti-correlation between a protein’s sequence and structure preference. Although this has been observed for single proteins, a systematic relationship between the two information contents has never been shown so far.

Finally, for several RBPs (particularly DGCR8, SFRS1, and YY1), our motif finder could recover different sequence motifs in different structural contexts. Existing approaches, such as *RNAcontext* and *GraphProt*, model the sequence and structure motif separately but do not give insights into which sequence motif is present in which structural context. Our approach is the first to extract this information and reveal different modes of RNA binding by a single RBP.

Second, our model is able to retrieve informative motifs and, once trained on a set of sequences, is easy to interpret. In fact, HMMs are graphical models and the learned sequence-structure motif can be easily visualized as a graph of state nodes with sequence logos. Given their high classification accuracy, *GraphProt* and *RNAcontext* are optimized for distinguishing real RBP binding sites from non-binding sites and detecting new binding sites rather than deriving a motif description for a certain protein of interest. Both tools train a classifier that provides an accurate protein-binding model, which can be applied to find RBP binding sites transcription-wise. Thanks to its graph-based feature encoding of the RNA structure, *GraphProt* takes into account a wider structural context around the RBP binding site and long-range interactions typically observed in RNA secondary structures. While this might help the tool in discriminating real binding sites from non-binding sites, the motif description derived from the classification model remains fuzzy. *MEMERIS* and ssHMM, in contrast, are motif finders designed for the extraction of the most prominent sequence and/or structure pattern from a dataset. Our analyses showed that the motifs recovered by the two motif finders possess a considerably larger information content than the motifs by *GraphProt* and *RNAcontext*. In many cases, the IC of the sequence motifs retrieved by *MEMERIS* is higher than the sequence-only IC of ssHMM. This is to be expected given that *MEMERIS* only optimizes the sequence motif model while ssHMM tries to optimize a joint sequence-structure motif. Both *MEMERIS* and ssHMM also achieved substantially higher motif recovery rates than the two classifiers on synthetic data.

Third, several evaluations confirmed that our motif finder correctly recovers motifs present in synthetic and biological datasets. Evaluation of our motif finder on synthetic data revealed its ability to retrieve 80% to 100% of the implanted motifs in all settings, in the absence of other confounding signals. This was also the case for motifs with very low sequence information content and fuzzy structural context. Significant differences in log-likelihoods between positive and negative test sequences demonstrated the good fit of the model to the protein’s binding preferences. Finally, analysis of CLIP-Seq datasets from RBPs with previously known binding motifs confirmed that the motif finder recovers correct motifs from biological data.

Fourth, ssHMM is faster than other tools. RNAs are flexible and dynamic molecules which can adopt multiple, stable secondary structures. Therefore, *RNAcontext* considers, for each nucleotide position, the probability over a certain number of possible structural contexts. This is time-consuming, as our runtime analysis showed, and prohibitive with a dataset of thousands of sequences. We circumvent this problem by making use of RNA shapes and Gibbs sampling to select the best shape, and thereby reducing notably the folding space of an RNA sequence, while still allowing different stable structures to be incorporated in the model training.

The runtime performance of our motif finder enables its application on large CLIPSeq datasets. While *MEMERIS* exhibits a quadratic runtime and *RNAcontext* is impeded by its slow structure prediction step, our approach is able to analyze even datasets with more than 20,000 sequences in reasonable time.

We have shown the ability of our method to recover motifs for few RBPs whose sequence-structure preferences are not fully characterized, such as DGCR8 and YY1. In the latter, ssHMM can also capture the mixture of two sequence motifs, one in the stem and the other one in a multiloop. Although not shown in this study, ssHMM can in principle be applied to any set of RNA sequences generated from experimental techniques other than CLIP-Seq, for example SELEX experiments. It requires only the FASTA file of the sequence set as input.

As future perspective, the motif models generated by ssHMM can be stored and used to search for motif hits in uncharacterized RNA sequences. For example, one could look for RBP motifs in long non-coding RNAs, a class of RNAs whose full spectrum of possible functions is still poorly understood. ssHMM can help annotating long non-coding RNA functions based on their most likely interaction partners.

## 4 Conclusions

We have developed a new efficient algorithm to determine the most probable sequence-structure motif, or combination of motifs, given a large set of RNA sequences. As RNA secondary structure is required for the molecular function of the RNA and for the specificity of RBP binding, and large-scale assays are becoming more and more popular in studying RNA-RBPs interactions, our method will contribute to the systematic understanding of such interactions.

## 5 Methods

In the following, we describe in detail the HMM that is used to represent the sequence-structure motif as well as the Gibbs sampling procedure for the estimation of the model’s parameters. We also explain how the synthetic and biological datasets were generated and how evaluations were performed.

### 5.1 The sequence-structure hidden Markov model

The sequence-structure hidden Markov model (ssHMM) models the RNA-protein binding site as a set of symbol-emitting states. The symbols are the four nucleotides A, C, G, and U. Each combination of binding site position *P* ϵ {1*..n*} and structural context *C* ϵ {*E, I, S, H, M*} is represented by exactly one state (Fig. 6). The states and transition probabilities of the HMM represent the different RNA structures and the transitions between them, respectively. The emissions and emission probabilities, on the other hand, represent the RNA nucleotides and the probabilities of them being observed in a specific structural context. The ssHMM can thus model the inter-dependencies between RNA sequence and structure preferences in RNA-protein binding. Particularly, it is able to model the occurrence of different sequence motifs in different structural context. The length *n* of the binding site motif to model needs to be chosen by the user. It is recommended to try different motif lengths for an RBP and to select the motif model which yields the highest probability.

**Figure 6:**
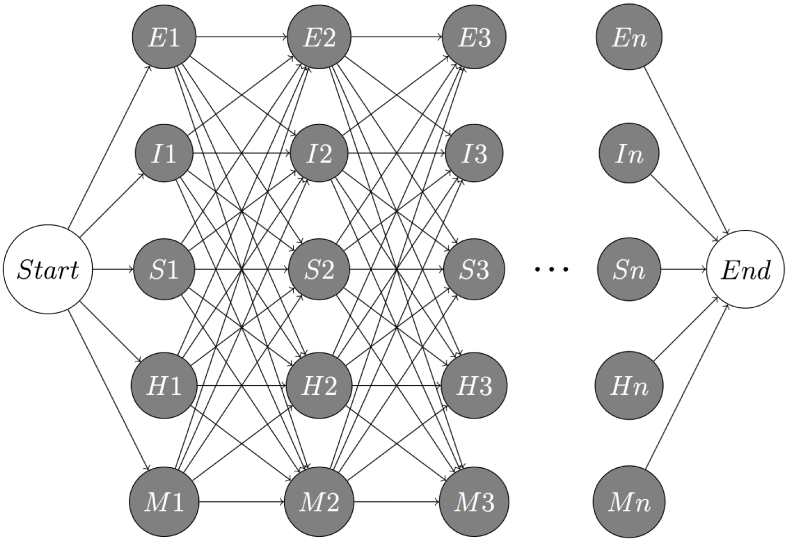
*Topology of the ssHMM - Each combination of binding site position (columns) and structural context (rows) is represented by one state. The structural contexts are E(xterior), I(nternal or bulge), S(tem), H(airpin), and M(ultiloop). Note that every transition in the model proceeds immediately to the next binding site position and that exactly n nucleotides are emitted by the HMM from Start to End.*

### 5.2 Model training

In order to train the ssHMM, the following two sets of variables need to be estimated:

#### Motif start positions

The ssHMM models solely an RBP’s binding motif, not entire RNA sequences. The binding motif is typically much shorter than the input RNA sequences and it is unknown where it is located in each of the long RNA sequences. This information, however, is needed to train the ssHMM (which models only the motif) with the long RNA sequences. As the motif length is given, it is sufficient to estimate the motif start position for every RNA sequence.

#### Best structure

It is hard to predict the secondary structure that an RNA adopts during the binding by an RBP solely based on thermodynamic calculations. RNAs are flexible chains of nucleotides which can often fold into multiple stable structures and whose structure can be influenced by other molecules. This is why most structure prediction tools compute several highly probable secondary structure conformations for each RNA nucleotide sequence. We employed the tools *RNAstructure* or *RNAshapes* [35, 36] to predict RNA structural states. *RNAstructure* predicts the lowest free energy structure as well as a number of suboptimal structures for a given RNA sequence. *RNAshapes*, in contrast, builds upon the concept of abstract shapes (i.e. classes of structures with similar features) to avoid predicting many highly similar and redundant structures. RNA structure prediction is a pre-processing step for our motif finder and the user can choose which tool to use in that step. Besides *RNAstructure* and *RNAshapes*, the user can compute secondary structures with any tool producing the required structure format. Regardless of the tool, the optimal structure, among several possible stable conformations, has to be chosen for every RNA sequence.

To estimate these two sets of unknowns we use a Gibbs sampling approach. We train the ssHMM while, at the same time, also estimating the two unknown variables for each sequence.

#### 5.2.1 Gibbs sampling procedure

At the outset of the Gibbs sampling, the unknown variables have to be initialized (Fig. 7), for instance by choosing random values. In each of the following iterations, one RNA nucleotide sequence together with its corresponding RNA structure sequences is held-out. An iteration consists of two estimation steps:

**Figure 7:**
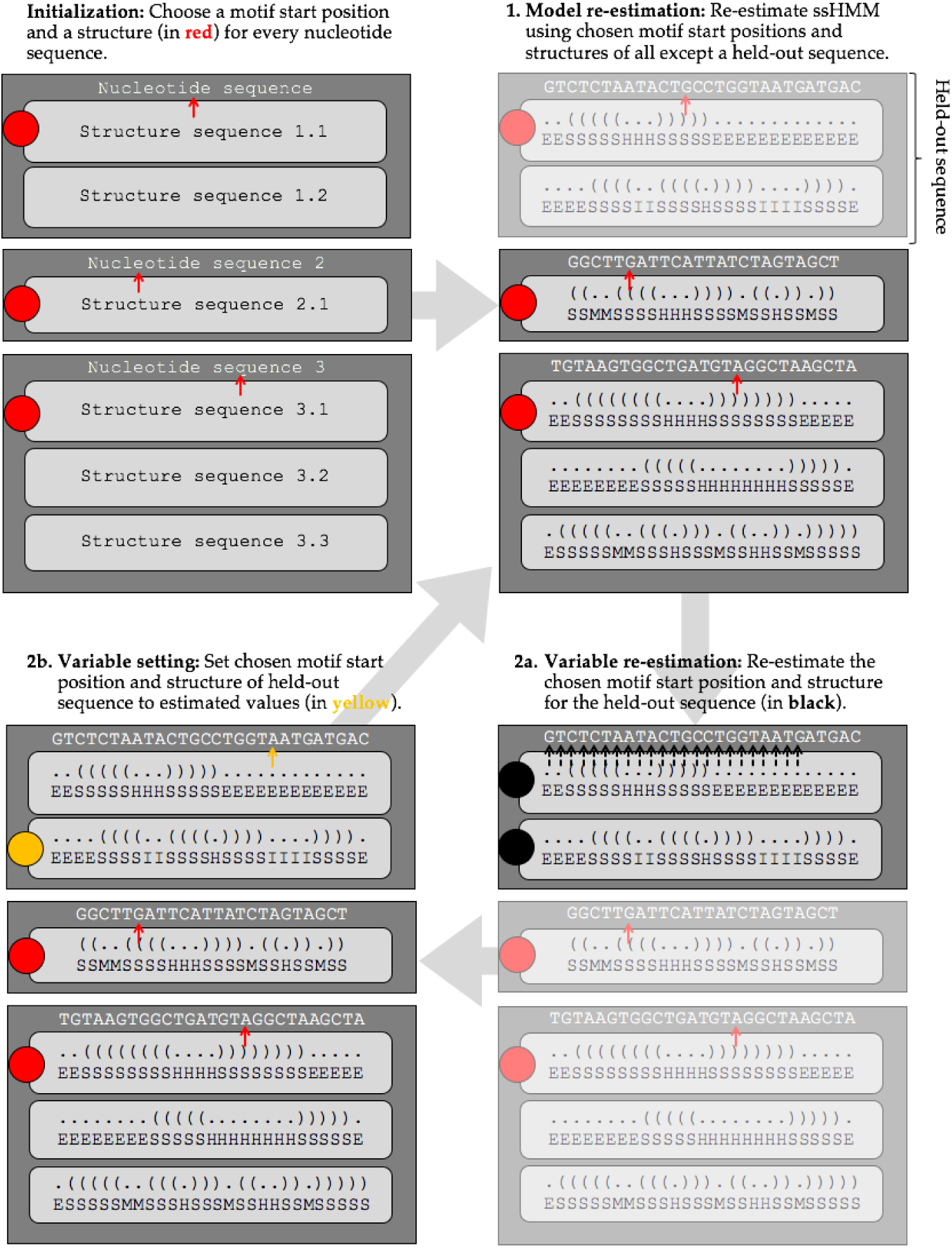
*Schematic overview of our Gibbs sampling approach - The dark gray boxes represent the RNA sequences. Each RNA sequence is characterized by its nucleotide sequence and several possible structure sequences (light gray). During initialization, a structure and a motif start (red) are set for each sequence. Every iteration consists of 3 steps: (1) One of the sequences is randomly chosen. Then, the ssHMM is re-estimated using all but this chosen sequence. (2) The two unknown variables of the held-out sequence are re-estimated. For this purpose, the probability of every combination of structure and motif start (black) given the ssHMM is calculated. (3) According to the distribution of these probabilities, a new structure and a new motif start is drawn (yellow). These three steps are repeated until termination.*

1. The ssHMM is re-estimated using all sequences except the held-out one. The current best structure and motif start positions can be used to retrieve both the nucleotide motif occurrence and the structure motif occurrence from each sequence. The nucleotide motif occurrence denotes a series of HMM emissions (an emission sequence) while the structure motif occurrence denotes a series of HMM states (a path). Using all state paths and emission sequences, it is possible to calculate a maximum likelihood estimate for the model parameters. To estimate the transition probabilities 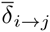, the number of transitions *t*_*i,j*_ between each pair of states *i* and *j* in the state paths can be counted. The maximum likelihood estimate of the transition probability between two states is then

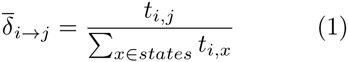 To estimate the emission probabilities 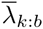, the number of emissions *e*_*k*_(*b*) of symbol *b* from state *k* can be counted in the emission sequences, and the maximum likelihood estimate of the emission probability from a state is then

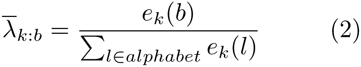 The initial probabilities 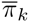 can be estimated by counting how often each state appears as the first state *s*_1_ in all state paths.
2. The motif start position and best structure of the held-out sequence is re-estimated given the ssHMM. For every possible combination of the two, we can calculate the conditional probability

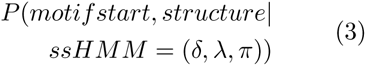

because motif start and best structure unambiguously define an emission sequence and a state path. We compute the joint probability of the emission sequence and the state path which is equivalent to the conditional probability. The new motif start position and best structure for the held-out sequence is drawn randomly according to the distribution of the conditional probabilities.

Thus, each iteration re-estimates the ssHMM and the unknown variables. After a large number of iterations, the algorithm converges on variables that maximize the likelihood of the ssHMM given the data. However, the likelihood improvement made in every iteration decreases over time. There is a point from which the improvement does not justify the required computational time anymore. The execution is therefore terminated when the improvement drops below a user-defined threshold (Additional File 1).

#### 5.2.2 Initialization

Gibbs samplers run the danger of becoming trapped in local optima. Therefore, it is advisable to choose these values carefully. In our approach, two initial values need to be determined at the start for every sequence: the best structure and the motif start position. From the structures, we always choose the one with the highest probability as determined by the structure prediction tool. For choosing the initial motif start positions, we implemented two different approaches:

##### Random

Motif start positions are drawn randomly. Depending on the drawn values, Gibbs sampling may yield very different results.

##### Baum-Welch

Motif start positions are determined using a sequence-only HMM. With the Baum-Welch algorithm, the sequence-only HMM is trained on the complete nucleotide sequences. Thus, the HMM is able to find the strongest sequence motif in the nucleotide sequences. Afterwards, the Viterbi algorithm is used to locate that sequence motif in each sequence. The starting index of the sequence motif is taken as the initial motif start position.

In a comprehensive analysis we performed, the Baum-Welch initialization approach yielded substantially better results than random initialization. Therefore, we used it to compare ssHMM against the other methods.

### 5.3 Dataset generation

For the evaluation of the motif finder, we collected two kinds of sequence datasets: randomly generated synthetic datasets and biological datasets derived from CLIP-Seq experiments on 18 different proteins.

#### 5.3.1 Synthetic datasets

Synthetic sequences are generated specifically to contain a certain implanted motif. For the generation of such sequences, we followed the protocol devised by Bahrami-Samani et al. but adapted it to our purposes [32]. The protocol describes the generation of datasets that contain a sequence motif of length 6. We generated twelve such datasets with different properties (Table 3). The properties were information content (1.0 / 0.5), background sequence type (uniform zeroth-order Markov chain / 3’UTR), and fraction of motifs in hairpin.

**Table 3:** *Properties of the twelve different synthetic datasets.*

| ID | Background seq. | Information cont. | Hairpin fraction |
| --- | --- | --- | --- |
| A | uniformly random | 1.0 | 100% |
| B | uniformly random | 1.0 | 50% |
| C | uniformly random | 1.0 | 10% |
| D | uniformly random | 0.5 | 100% |
| E | uniformly random | 0.5 | 50% |
| F | uniformly random | 0.5 | 10% |
| G | 3' UTR | 1.0 | 100% |
| H | 3' UTR | 1.0 | 50% |
| I | 3' UTR | 1.0 | 10% |
| K | 3' UTR | 0.5 | 100% |
| L | 3' UTR | 0.5 | 50% |
| M | 3' UTR | 0.5 | 10% |

Each of these twelve datasets is comprised of 100 sequence sets. A sequence set consists of 2,000 RNA sequences with their corresponding shapes and structures. For each sequence set, one random position probability matrix (PPM) of length 6 with a given average information content was created and stored for later evaluation.

For each of the 2,000 sequences in the sequence set, a motif occurrence was drawn from the PPM. This motif occurrence was implanted into a background sequence of length 50 at a random location. Depending on the concrete dataset, the background sequence was either generated by drawing from a uniform distribution over the four nucleotides (datasets A to F) or randomly sampled from the set of human three prime untranslated regions (3’UTRs, datasets G to M). 3’UTRs are genomic regions known to contain many RBP binding sites. They are therefore well-suited to yield realistic background sequences for the synthetic datasets.

Depending on the concrete dataset, it was ensured that a certain fraction of the 2,000 motifs is implanted into a hairpin loop. We wanted to assess the influence of the structural context on the motif recovery. In order to deliberately implant a motif in a hairpin loop, we inserted the reverse complement of the sequence located next to the motif onto the motif’s other side. Very often, the resulting sequence will fold such that the motif lies in a hairpin loop. To confirm this, we ran the structure prediction tool on the resulting sequence and retained the sequence only if the entire motif was predicted to be in a hairpin.

#### 5.3.2 CLIP-Seq datasets

We retrieved 21 different CLIP-Seq datasets for 18 different RBPs from various sources (Additional File 1). Most datasets were downloaded from the *doRiNA*, a database of manually curated RBP binding site data [51]. With the exception of two mouse datasets, all experiments were conducted in human HEK293 and HeLa cells. From the 21 datasets, 14 were generated with PAR-CLIP, 6 with HITS-CLIP, and 1 with iCLIP and provided as genomic coordinate files in Browser Extensible Data (BED) file format. All CLIP-Seq datasets are regarded as positive datasets because they contain the RBP binding sites.

Both *RNAcontext* and *GraphProt* require positive as well as negative (i.e. bound and unbound) sequences to train on. Therefore, we created a corresponding negative dataset for each of the 21 positive CLIP-Seq datasets by randomly drawing regions from the genome whose lengths match those of the positive binding sites. Thus, we obtained datasets that contain the same number of sites as their positive counterparts and whose sites have the same lengths. As a result of the random drawing procedure, we expect the vast majority of these sites to be located outside of real biological binding sites (i.e. to be negatives).

Starting from the positive BED file, the following pre-processing steps were taken to obtain the final sequence files:

1. Filter positive BED file (remove low-scoring regions and regions shorter than 8nt or longer than 75nt)
2. Generate negative BED file (draw random regions from genome)
3. Elongate both BED files (by 20 bases on each side)
4. Fetch genomic sequences for elongated BED files to obtain sequence files
5. Transform sequence files into viewpoint format (i.e. original regions in uppercase, elongations in lowercase)

For both types of datasets, secondary structures were predicted with either *RNAshapes* or *RNAstructure*. For the evaluations, we exclusively used *RNAshapes* because its predictions are less redundant. *RNAshapes* was run with command line options -o 1 (choosing output type 1) and -r (calculates structure probabilities). Otherwise, default parameters were used. *RNAshapes* produces output in the dotbracket format which consists of dots (representing unpaired nucleotides) and matching brackets (representing base pairs). The dotbracket output was converted to a string of structural contexts using the *forgi* 0.2 python library [52]. The string encodes the predicted structural context of each nucleotide in the input sequence with a symbol. The symbols are E for exterior loop, I for internal loop, S for stem, H for hairpin loop, and M for multiloop.

The secondary structures predicted by *RNAshapes* were only used to train our motif finder. Although *GraphProt* also uses *RNAshapes*, structure prediction is already integrated into *GraphProt*’s training process. *RNAcontext* and *MEMERIS* use *Sfold* and *RNAfold*, respectively.

### 5.4 Evaluation method on synthetic datasets

The motif recovery performance of the motif finders on synthetic datasets was evaluated in terms of the Kullback-Leibler (KL) divergence, applied here to measure the difference between two position probability matrices. The KL divergence between two probability distributions P and Q over the four nucleotides is defined as:

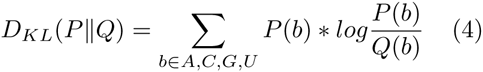

After training on the synthetic sequences, all motif finders produce a sequence logo that can be expressed in terms of a PPM. This PPM contains the probability of every base at every position and we call it the *recovered PPM*. The PPM that was used to generate the synthetic sequences is called the *original PPM*. The KL divergence between these two PPMs expresses how well the motif finder recovered the motif from the sequences. We used an absolute threshold of 1.0 to classify recovered motifs as either ‘successfully recovered’ or ‘not recovered’.

### 5.5 Evaluation method on CLIPSeq datasets

To evaluate the performance of our motif finder on all 21 CLIP-Seq datasets, we first checked that the trained motif finder can distinguish between real binding sites (positives) and background sites (negatives). Secondly, we carried out a qualitative analysis of the recovered motifs, assessed their resemblance to motifs from the literature and interpreted the biology behind some of the motifs.

To assess whether ssHMM can distinguish between real binding sites and background sites, we first trained it on a positive training set of putative RBP binding sites. The training set was compiled by randomly sampling 90% of all positive sequences in the CLIP-Seq dataset. The remaining 10% were reserved for the positive test set. A negative test set was compiled by randomly sampling a number of negative sequences equal to the number of sequences in the positive test set. We computed the log-likelihoods of the sequences from the separate positive and negative test sets. Then, we applied the non-parametric Wilcoxon rank sum test to test for significant differences between log-likelihoods of positive and negative test sequences. For a p-value smaller than 0.01, we regarded the difference to be significant. All training and test data can be found on Github at https://github.molgen.mpg.de/heller/ssHMM_data.

### 5.6 Calculation and comparison of motif information content

Depending on the underlying alphabet *A*, the information content of a binding motif position can range from 0 to log_2_|*A*|. Consequently, the maximum information content per position of a nucleotide sequence motif is log_2_ 4 = 2. The maximum information content per position of a structural motif with *|A|* = 5 is log_2_ 5 ≈ 2.32 and of a sequence-structure motif it is log_2_(4 * 5) ≈ 4.32. To calculate the information content of a motif position, the frequency *f*_*s*_ of each symbol *s ∈ A* is required (e.g. from a PPM) [37]. Then, the Shannon entropy *H* and small-sample correction *e* of that position are defined as

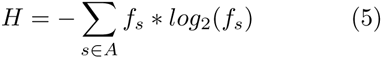

and

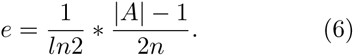

Finally, the information content of that position can be computed as [37]

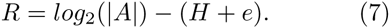

The total information content of a motif can be obtained by summing over the information content of all motif positions. To ensure a fair comparison of the information contents, one has to distinguish two ways of calculating the information contents: 1) Information content from the top sequences, and 2) Information content directly from the model.

#### 5.6.1 Information content from the top sequences (comparison with GraphProt)

*GraphProt* computes sequence and structure logos from the 1,000 highest-scoring k-mer nucleotide sequences and structure profiles. From these 1,000 sequences, the frequency *f*_*s*_ of each symbol *s* ∈ *A* in each motif position can be calculated by counting. These frequencies serve as input to compute the information content as described above.

To ensure comparability, we followed a similar procedure to obtain the information content of sequence motifs produced by our motif finder. We calculated the information content on the 1,000 sequences with the best score given our trained motif model. First, the estimated motif start and best structure of each of these sequences was obtained from the trained model. They pointed directly to each sequence’s motif occurrence which made it possible to align the 1,000 motif occurrences (sequence and structure separately). Then, the frequencies were counted and the three different types of information content (sequence, structure and combined sequence-structure) were computed.

#### 5.6.2 Information content directly from the model (comparison with RNAcontext)

*RNAcontext* directly produces a PPM representing its inner model. To compare against *RNAcontext*, we obtained a second set of information contents directly from the ssHMM. First, a sequence motif (or PPM) was extracted from the trained ssHMM by averaging over all paths in the model. The average was weighted by the transition probabilities so that more likely structural contexts have a bigger impact on the sequence motif than less likely contexts. Then, the information content was computed.

The information contents calculated from the top sequences are naturally larger than those directly from the model. The reason is that those sequences, which are most likely given the model, are more homogeneous than the set of all sequences. Consequently, the resulting motif is more clearly defined.

### 5.7 Software libraries

HMMs have been implemented by numerous software packages in several programming languages. For this project, we built upon the *general hidden Markov model* (GHMM) library [53]. It is written in C but provides wrappers for interactive use in Python and R. It implements all basic and many advanced aspects of HMMs, such as multivariate Gaussian mixture HMMs and pair HMMs.

### 5.8 Program parameters

All three analyzed tools as well as our method have multiple parameters that heavily influence their performance. Below, we describe how we chose the best parameters for each tool. The full parameters are shown in the Additional File

#### 5.8.1 MEMERIS

*MEMERIS* inherits most parameters from *MEME* and adds one of its own: pi, which specifies the pseudocount that is used to flatten the prior probability distribution of the motif starts. In essence, it defines how strongly *MEMERIS* prefers motifs in a single-stranded context. The lower its value, the more does it prefer single-stranded motifs. A very high pseudocount of 10 or higher results in a *MEME*-like behavior which ignores structure completely. We ran *MEMERIS* with four different values for pi: 0, 1, 5, and 20.

Besides that, we left most parameters at their default values. Motif length was naturally set to 6. A uniform background distribution was assumed because the sequences are too short to reliably estimate the background distribution of the characters from them. Lastly, the distribution of motif sites was set to OOPS which means that *MEMERIS* expects exactly one motif occurrence per sequence.

#### 5.8.2 RNAcontext

*RNAcontext* has two important parameters. w specifies the range of motif lengths. A range of 4-7, for instance, means that the algorithm searches for motifs starting from length 4 until length 7. *RNAcontext*’s initialization procedure uses previously learned models for smaller motif lengths to initialize longer motif lengths. Therefore, they suggest to use, e.g. 4-6 instead of 6-6 when looking for motifs of length 6. We found, however, that w set to 6-6 performed much better than 4-6 which is why we use that setting. Depending on the initialization, *RNAcontext* can generate different results. The parameter *s* defines the number of different initializations that are tried to obtain the best result. For this parameter, we use the default value 5.

#### 5.8.3 GraphProt

*GraphProt* has eight parameters which can be optimized in a dedicated parameter optimization step (program option -ls). We ran parameter optimization on the first sequence set of each dataset and used these optimized parameters for the entire dataset.

#### 5.8.4 Our motif finder

Our motif finder has six different parameters which are explained in more detail in Additional File 1:

- Motif length
- Initialization
- Block size
- Flexibility
- Termination interval
- Termination threshold

In Additional File 1, we also explain how we chose the best-performing values for these parameters: motif length = 6, initialization = Baum-Welch, block size = 1, flexibility = 0, termination interval = 100, termination threshold = 10.

## 6 Data access

The ssHMM software is available for download at https://github.molgen.mpg.de/heller/ssHMM. It contains three main command line scripts. New CLIP-Seq datasets can be pre-processed for analysis with the *preprocess dataset* script. Subsequently, the datasets can be analyzed with the *train seqstructhmm* or *batch seqstructhmm* scripts. Synthetic and CLIP-Seq datasets used in this study can be found at https://github.molgen.mpg.de/heller/ssHMM_data.

## 8 Author’s contributions

DH contributed to conceive the study, collected the data, implemented the method and performed all experiments. RK provided help with the method’s implementation, analysis and supervision of the study. UO provided help with the evaluation of the model and valuable expertise on motif identification algorithms. MV contributed to the algorithm’s idea and provided valuable help in the in the interpretation of the results. AM conceived the idea and supervised all the steps of the study. DH and AM wrote the manuscript. RK and MV commented on the manuscript at all steps. All authors have read and approved the manuscript for publication.

## 9 Acknowledgements

The authors would like to thank the developers of the GHMM python package that is used by ssHMM. The project was supported by the Freie Universit¨at Berlin within the Excellence Initiative of the German Research Foundation (DFG). Support by the SFB-TR 84 is also gratefully acknowledged.

## 10 Additional Files

### 10.1 Additional File 1 — Supplementary information

This file contains supplementary information about the ssHMM (parameters, motif output) and the evaluation method. Additionally, it contains all P-values from the Wilcoxon rank sum test and results of the runtime comparison.

### 10.2 Additional File 2 — Full CLIP-Seq results

This file contains a table with the recovered motifs for all 21 CLIP-Seq datasets, including results by ssHMM, MEMERIS, RNAcontext, and GraphProt (with motif information contents).

### 10.3 Additional File 3 — ssHMM motifs for RNAshapes and RNAstructure

This file contains a table with the ssHMM motifs for all 21 CLIP-Seq datasets based on structures predicted by RNAshapes or RNAstructure.

